# Effect of fluid control on the affective state of laboratory macaques

**DOI:** 10.1101/2025.10.10.681594

**Authors:** Janire Castellano Bueno, Alexandra Paraskevopoulou, Christopher W. Miller, Christopher I Petkov, Michael C Schmid, Alexander Thiele, Melissa Bateson, Colline Poirier

## Abstract

Fluid control protocols are widely used in neuroscience to motivate laboratory macaques to engage with behavioural tasks. Despite strong evidence that the physiology of the animals is not compromised by such protocols, fluid control remains controversial due to its potential impact on the psychological well-being of the animals. To address this concern, we investigated the effect of fluid control on the affective state of 23 socially-housed adult macaques (10 females) engaged in neuroscience experiments. The protocol involved up to five consecutive days of fluid control per week, followed by a minimum of two days with unrestricted fluid access. The affective state of the animals was primarily assessed by quantifying the frequency of pharmacologically-validated behavioural indicators of high-arousal negative affect (self-scratching, body shaking, self-grooming). The analysis was subsequently extended to validated behavioural indicators of low-arousal negative affect (*Inactive not alert*) and other behaviours suspected of indicate high-arousal negative affect but lacking proper validation (pacing, yawning). In total, 700 hours of video footage spanning up to seven years of intermittent fluid control per animal were analysed. Despite this extensive dataset, the study found no significant impact of fluid control on average, or any evidence of habituation or sensitization over the years on any of the affective state indicators. Additional results indicate that these null results are not due to a lack of sensitivity, supporting the view that fluid control, as implemented in this study, does not have an adverse impact on the psychological well-being of laboratory macaques. We argue that macaque welfare will be best served by focusing future refinement on other procedures.

## Introduction

Macaques are capable of performing complex cognitive and behavioural tasks requiring high visual acuity, manual dexterity and behavioural flexibility. Together with their phylogenetic proximity to humans, these abilities make them valuable animal models in neuroscience research. To obtain scientifically reliable results, many behavioural tasks implemented in neuroscience experiments require a large number of consecutive trials, reached only when macaques are highly motivated to perform the task. To achieve such motivation levels, neuroscientists generally use fluid or food control. These methods consist of not providing any fluid or food before the experimental session and using small amounts of fluid or food during the session as rewards (the amount of fluid or food provided to the animals after the session depends on the specific protocol). Depending on the nature of the task and methodological constraints, fluid control is typically preferred when: (1) the experimental sessions require an extensive number of small rewards to be delivered; (2) the reward needs to be automatically delivered in a controlled and standardized way; and/or (3) eating-related chewing movements would induce artefacts in the collected data (e.g., electrophysiological and MRI recordings).

The effect of fluid control on the physical health of laboratory macaques has been assessed by several research groups implementing slightly different fluid control protocols (Yamada et al., 2010; Gray et al., 2016; Wegener et al., 2021). In these studies, fluid control has been consistently found to have no adverse impact on blood hydration levels or kidney function. However, the potential impact of such protocols on the psychological well-being of macaques has been debated since the nineties (Orlans, 1991; Desimone et al., 1992; Prescott et al., 2010; Westlund, 2012), due to a lack of systematic studies. Three relatively recent studies have tried to fill this gap but with inconclusive results. Two studies implemented a behavioural approach to assess the affective state of the animals and found no indication of negative impact (Hage *et al*., 2014; Gray *et al*., 2016). However, both studies had small sample sizes, (four and seven individuals, respectively), and only male subjects. Most importantly, the behavioural categories selected as welfare indicators lacked validation as indicators of negative affective states and may have been insensitive to small changes in affect. A third study used salivary cortisol as an affective state indicator and found evidence for increased cortisol levels in 16 males (Pfefferle *et al*., 2018). However, the increase was very small compared to natural circadian variation in cortisol levels, and the interpretation of this result is questionable, given the body of literature showing that increased cortisol indicates high arousal and can be indicative of both negative and positive affective states (Ralph and Tilbrook, 2016). An alternative interpretation of the observed increased cortisol is that animals experienced a high-arousal positive affective state, driven by the expectation of cognitively-challenging tasks, a level of stimulation unrivalled by the cage environment despite the presence of enrichment. An additional concern is that cortisol levels are insensitive to negative affective states associated with low arousal. On top of these limitations, none of the previous studies has considered potential habituation or sensitisation to fluid control protocols over time. Considering that macaques involved in neuroscience research can be subjected to fluid control protocols for several years, investigating the longitudinal effect of these protocols is critical for understanding the welfare of these animals.

The present study aimed to identify the acute effect of fluid control on the affective state of a large (15+) sample of female and male laboratory macaques and to investigate whether this effect varied over the years and with the number of consecutive days of fluid control and the severity of fluid control (expressed as the amount of fluid divided by the animal weight). Using a quasi-experimental design, whereby neuroscientists decided whether or not each animal was fluid controlled on any given day according to their experimental needs, we tested the effect of fluid-control on behavioural indicators of affective state. We hypothesised that if fluid control had an impact on the affective state of macaques, it would be similar to the effect of thirst in humans, namely a negative affective state associated with high arousal (McKinley, 2009). Affective states were thus primarily assessed using pharmacologically-validated behavioural indicators of such affective state, namely displacement behaviours combining self-scratching, self-grooming and body shaking (Maestripieri *et al*., 1992; Schino *et al*., 1996). In case of null results, secondary analyses were planned using behaviours suggested to be indicative of high-arousal negative affective states but not yet validated, namely yawning and stereotypic pacing (Lagarde *et al*., 1990; Major *et al*., 2009; Poirier and Bateson, 2017), as well as a pharmacologically-validated indicator of negative affective state associated with low arousal, namely *Inactive not alert* behaviour (Jackowski *et al*., 2011; Qin *et al*., 2015). We predicted that if fluid control has an adverse effect on affective state, the frequency of the target behaviour should be higher when the animal had been fluid controlled compared to when it had received fluids *ad libitum*), either (1) overall, or (2) at least in the first or last instances of fluid control (in case of respectively habituation or sensitization), and/or (3) after the highest number of consecutive days of fluid control and/or (4) after the most severe cases of fluid control.

## Results

### Descriptive statistics

Home-cage spontaneous behaviour data were recorded twice a year between 2014 and 2023, resulting in 18 periods of data collection subsequently referred to as time points. Time points typically took place during spring and autumn and lasted 5 to 8 weeks. During these periods, data were recorded every day except on Saturdays and Sundays, as human presence and husbandry routines are significantly different at the weekend compared to weekdays. For each daily session, the identity of the subject, the date and the corresponding time point were encoded, the frequency of each target behaviour was computed, and fluid control data were retrieved from historical records. We extracted whether the animal had been fluid controlled or not (*Condition* variable), the relative time since its first ever fluid control (*Years* variable), the number of consecutive days the animal had been fluid controlled (*Duration* variable), the amount of fluid drank (*Fluid amount* variable) and the weight of the animal (*Weight* variable). The distribution of the data is illustrated in Figure 1.

**Figure 1.**
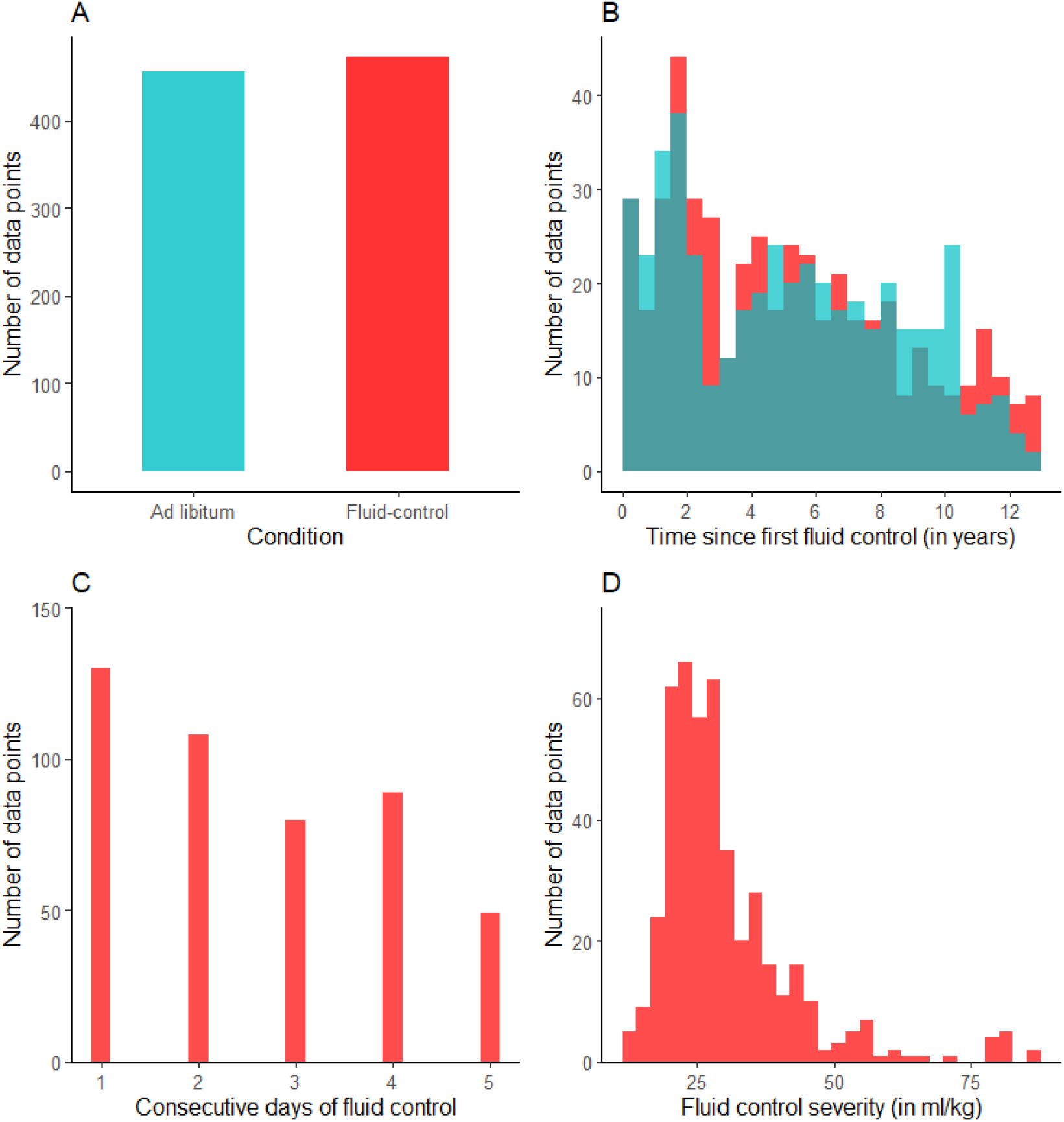
Data distribution. A: Total number of data points per condition (*Ad libitum* vs. Fluid control). B. Distribution of ad-libitum data (in blue) and fluid-controlled data (in red) over time (expressed in years since first fluid control). C: Distribution of fluid-controlled data according to fluid control duration, expressed as the number of consecutive days of fluid control at the time of the behavioural measurement. D: Distribution of fluid-controlled data according to fluid control severity (defined as ml of fluid given per kg body weight). The total sample size was 928 datapoints (including 472 in the fluid-control condition), coming from 23 individuals (10 females).

Data were modelled using general linear mixed models. To control for the non-independence of data, the variables *Subject* and *Time point* were declared as random effects in all models, with *Time point* nested within *Subject*. Sources of variation in the data were separated into within- and between-subjects effects by centring independent variables within each subject (van de Pol and Wright, 2009). The dependent variable was the frequency of the affective state indicator, while independent variables declared as fixed effects varied according to the specific model. To test our main hypothesis (i.e. effect of fluid control on the frequency of the behaviour), Model 1 included the independent variables *Condition* and *Years*, as well as their interaction in order to test for potential habituation or sensitization. To test the hypothesis that the frequency of the behaviour increases with the number of consecutive days of fluid control and/or with its severity, Model 2 was restricted to data points when the subjects were fluid controlled and included the independent variables *Duration, Fluid amoun*t, *Weight* and *Years* (since using ratios as independent variables does not allow determination of whether the effect is driven by the numerator or the denominator, severity, defined as *Fluid amount*/*Weight*, was modelled using both variables, with *Fluid amount* the variable of interest and *Weight*, used as a control variable). In case of a significant effect of *Duration* and/or *Fluid amount*, we subsequently tested the hypothesis that the frequency of the behaviour was higher in the most extreme cases of fluid control (defined for duration as 5 consecutive days, and for severity as below 20 ml/kg) compared to when individuals had *ad libitum* access to fluid (Model 3). This last analysis was restricted to subjects and time points where data were available from each condition and included the independent variables *Condition* and *Years*. For each model, assumptions from the general linear model were tested and data transformed if necessary. A square root transformation was applied to the following affective state indicators: yawning, pacing and *Inactive not alert*. For the primary analyses, the significance threshold was set at 0.05. For the secondary analyses the threshold was adjusted for multiple tests, using the Bonferroni correction (Bonferroni, 1936) and set to 0.016.

### Primary analyses: Displacement behaviours

Model 1 revealed no significant effect of *Condition*, with an average frequency of displacement behaviours of 40.2 % when subjects had *ad libitum* access to fluids and 40.7 % when they were fluid controlled (at an average time of 4.3 years after first fluid control). There was a significant effect of *Years*, with the frequency of displacement behaviours decreasing over the years, but no significant interaction between *Condition* and *Years*, indicating no support for habituation or sensitization effect (Fig. 2A, Table 1).

**Table 1.**
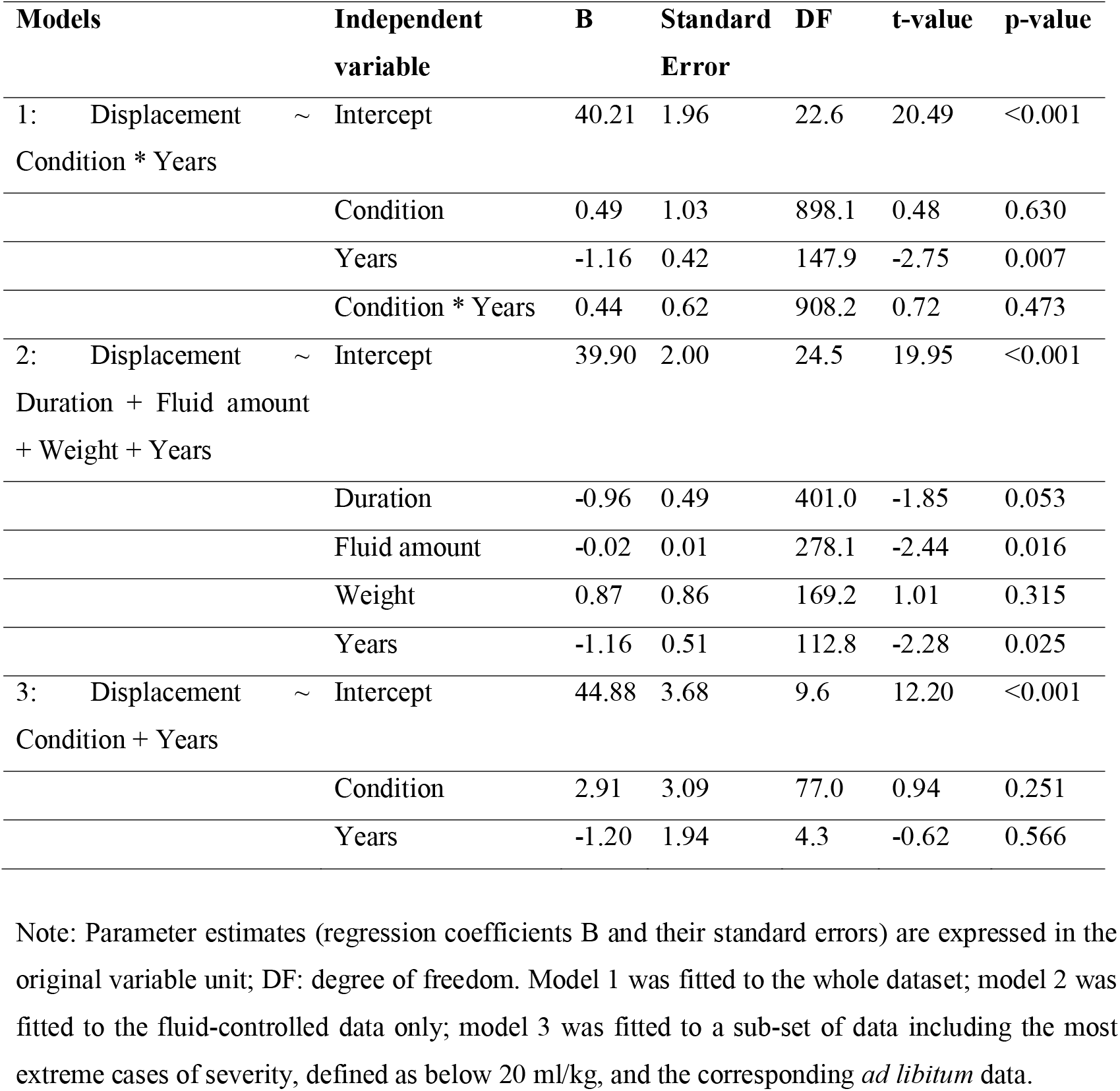
Statistical results for Displacement behaviours.

**Figure 2.**
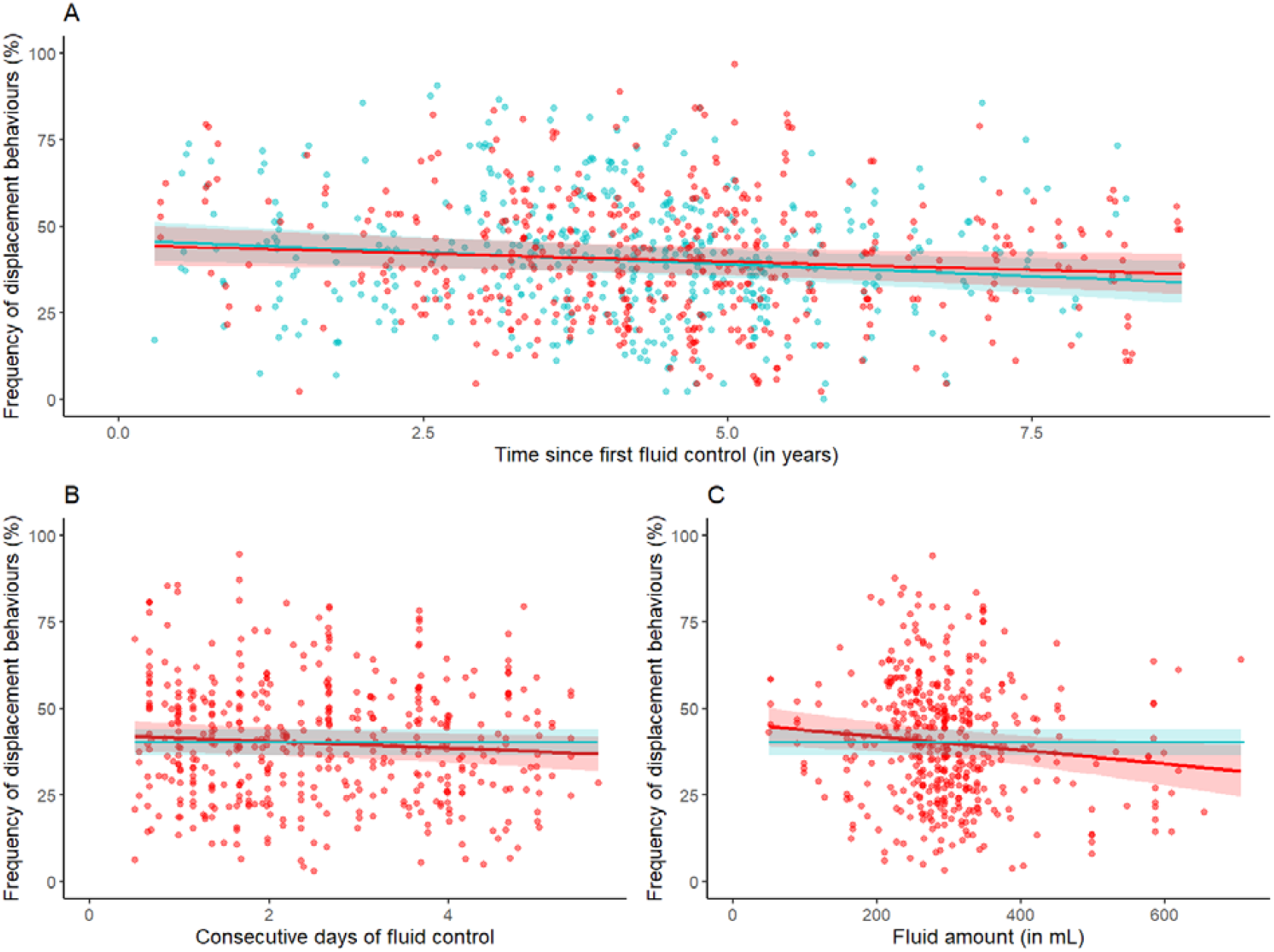
Changes in the frequency of displacement behaviours. (A) Interactive effect of condition (fluid-controlled vs *ad libitum*) with time since first fluid control (in years) (B) Effect of fluid control duration (expressed in number of consecutive days of fluid control). (C) Effect of fluid amount (expressed in ml). Blue: *Ad libitum*; Red: Fluid control. Dots correspond to mean-centred data, adjusted for the effect of other covariates when present in the models (i.e., *Fluid amount, Weight* and *Years* in panel B and *Duration* and *Years* in panel C), with the mean value of *Years* (A), *Duration* (B) and *Fluid amount* (C) added to make the x-axes meaningful. The lines and their ribbons correspond to the linear fit of the data and their 95% confidence interval, after accounting for random effects. For comparison purposes, the estimated frequency of the behaviour when subjects had *ad libitum* access to fluids (intercept of model 1b) and its 95% confidence interval were superimposed in panels B and C (blue lines and their ribbon).

Model 2 revealed no significant effect of *Duration*, but a significant effect of *Fluid amount*, with the frequency of displacement behaviours decreasing with fluid amount (while controlling for *Weight*), and thus increasing with severity (Table 1, model 2; Fig 2, panels B and C). Superimposing the estimated frequency of displacement behaviour and its confidence interval when individuals had a*d libitum* access to fluid (intercept from Model 1) revealed a large overlap of the confidence intervals (left-hand side of panel C in Fig. 2), suggesting that despite a higher frequency of the behaviour when macaques were given small amount of fluid (compared to larger amounts), this frequency did not become significantly higher that when animals had *ad libitum* access to fluids. This hypothesis was formally tested with model 3, which revealed no significant effect of condition in the most severe cases of fluid control (Table 1, model 3).

### Secondary analyses: Yawning

Fitting data with Model 1 revealed no significant effect of *Condition*, with an average yawning frequency of 2.3 % when subjects had *ad libitum* access to water and 2.2 % when they were fluid controlled. It also showed no significant effect of *Years* and no significant interaction, indicating no support for habituation or sensitization effect (Fig. 3A, Table 2). Model 2 revealed no significant effect of *Fluid amount* or *Duration* (Fig. 3B and 3C). (Please note that while the *Duration* effect would have been deemed significant without the Bonferroni correction, it is in the opposite direction as the one expected: yawning frequency decreased with longer duration of fluid control.)

**Table 2.**
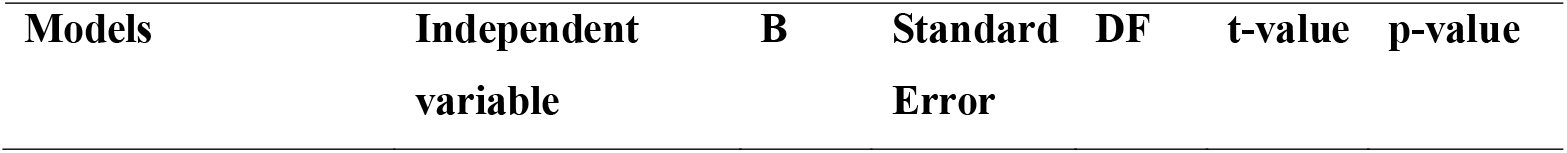
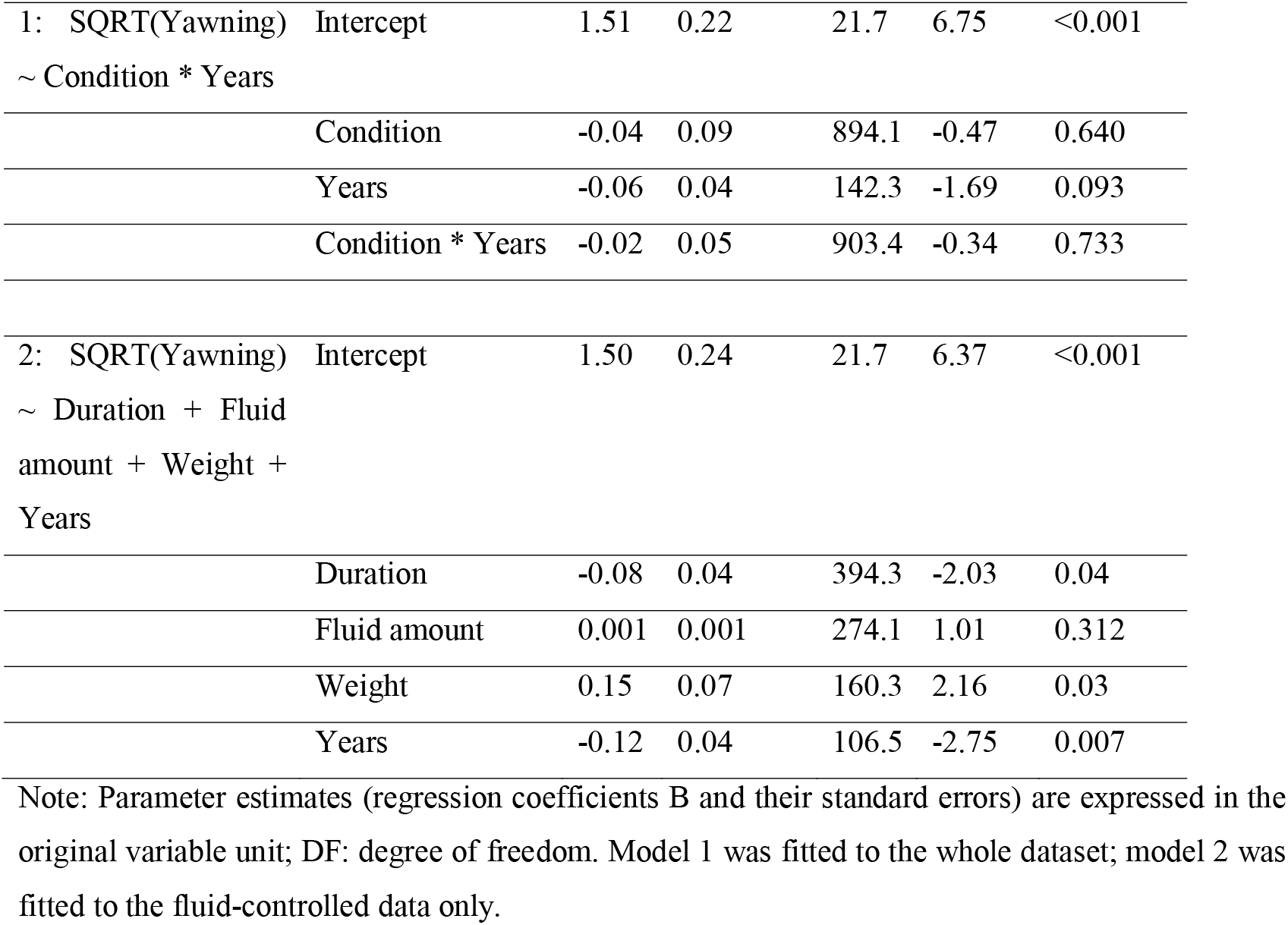
Statistical results for yawning behaviour, after square root transformation.

**Figure 3.**
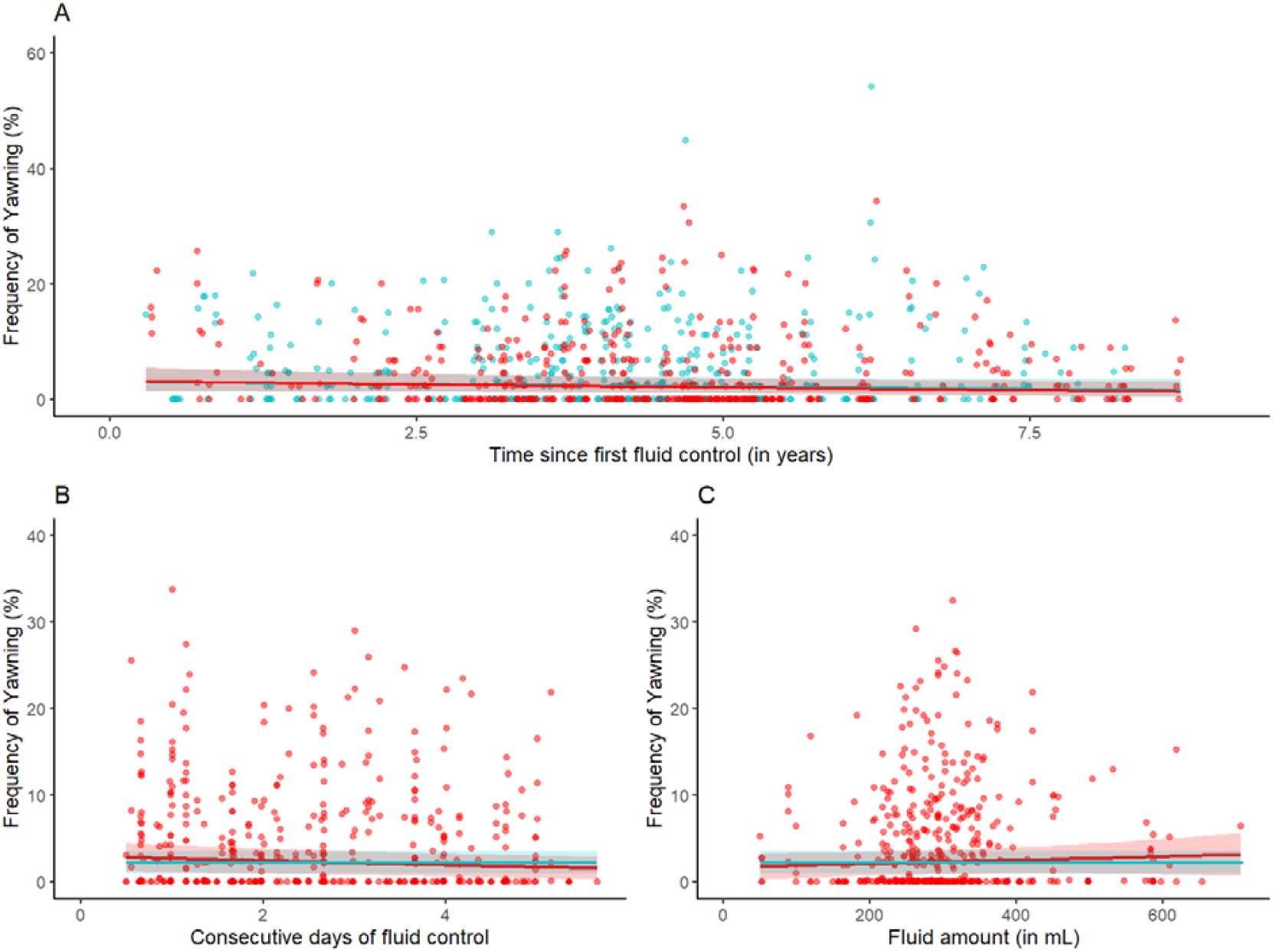
Changes in the frequency of yawning behaviour. (A) Interactive effect of condition (fluid-controlled vs *ad libitum*) with time since first fluid control (in years) (B) Effect of fluid control duration (expressed in number of consecutive days of fluid control). (C) Effect of fluid amount (expressed in ml). Blue: *Ad libitum*; Red: Fluid control. For more details, see legend of Figure 2.

### Secondary analyses: Pacing

For the analysis of pacing behaviour, the dataset was restricted to animals that displayed pacing behaviour at least once. As a result, the main dataset was reduced to 668 sessions from 16 subjects (5 females). Fitting data with Model 1 revealed no significant effect of condition, with an average pacing frequency of 4.7 % when subjects had a*d libitum* access to water and 5.6 % when they were fluid controlled). It also showed a significant effect of time, with the frequency of yawning decreasing over time but no significant interaction (Fig. 4A, Table 3). Model 2 revealed a significant effect of duration in the opposite direction than what was expected in case of negative impact of fluid control and no significant effect of fluid amount (Fig. 4B and 4C).

**Table 3.** Statistical results for Pacing behaviour, after square root transformation.

| Models | Independent variable | B | Standard Error | DF | t-value | p-value |
| --- | --- | --- | --- | --- | --- | --- |
| 1: SQRT(Pacing) ~ Condition * Years | Intercept | 2.16 | 0.48 | 14.7 | 4.51 | <0.001 |
|  | Condition | 0.20 | 0.15 | 612.3 | 1.32 | 0.186 |
|  | Time | -0.41 | 0.08 | 104.1 | -4.92 | <0.001 |
|  | Condition * Years | 0.02 | 0.09 | 624.9 | 0.21 | 0.837 |
| 2: SQRT(Pacing) ~ Duration + Fluid | Intercept | 1.98 | 0.48 | 14.5 | 4.14 | <0.001 |

| amount + Weight +<br>Years |  |  |  |  |  |  |
| --- | --- | --- | --- | --- | --- | --- |
|  | Duration | -0.22 | 0.07 | 258.5 | -3.20 | 0.002 |
|  | Fluid amount | 0.001 | 0.001 | 262.5 | 0.49 | 0.628 |
|  | Weight | 0.50 | 0.13 | 138.8 | -3.76 | <0.001 |
|  | Years | -0.24 | 0.09 | 79.0 | -2.54 | 0.013 |

Note: Parameter estimates (regression coefficients B and their standard errors) are expressed in the original variable unit; DF: degree of freedom. Model 1 was fitted to the whole dataset; model 2 was fitted to the fluid-controlled data only.

**Figure 4.**
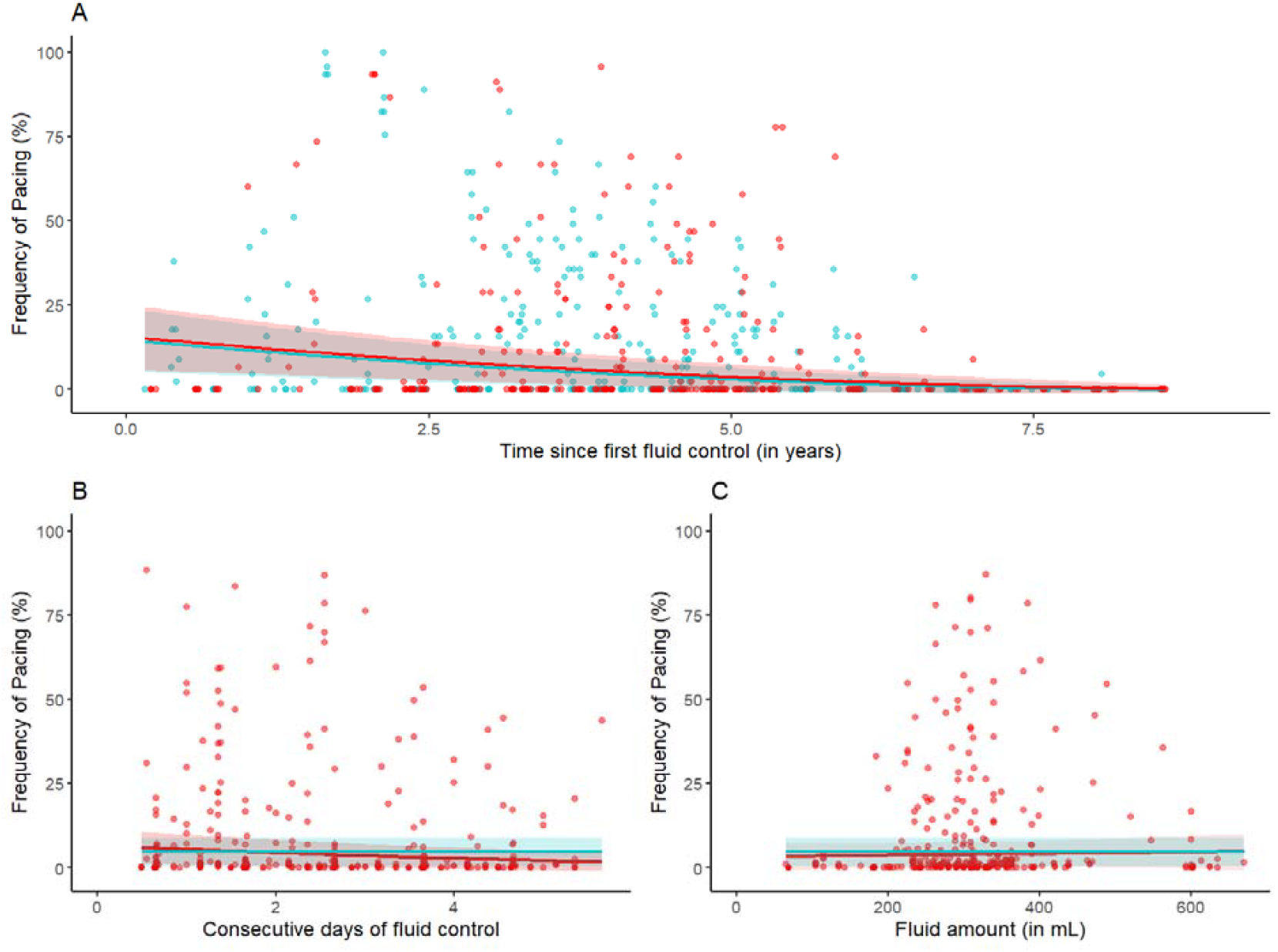
Changes in the frequency of Pacing behaviour. (A) Interactive effect of condition (fluid-controlled vs *ad libitum*) with time since first fluid control (in years) (B) Effect of fluid control duration (expressed in number of consecutive days of fluid control). (C) Effect of fluid amount (expressed in ml). Blue: *Ad libitum*; Red: Fluid control. For more details, see legend of Figure 1.

### Secondary analyses: *Inactive not alert*

Fitting data with Models 1 revealed that no significant effect of condition, with an average frequency of *Inactive not alert* of 2.3 % when subjects had *ad libitum* access to water and 2.5 % when they were fluid controlled. It also showed a significant effect of time, with the frequency of *Inactive not alert* increasing over time, but no significant interaction (Fig. 5A, Table 4).

**Table 4.**
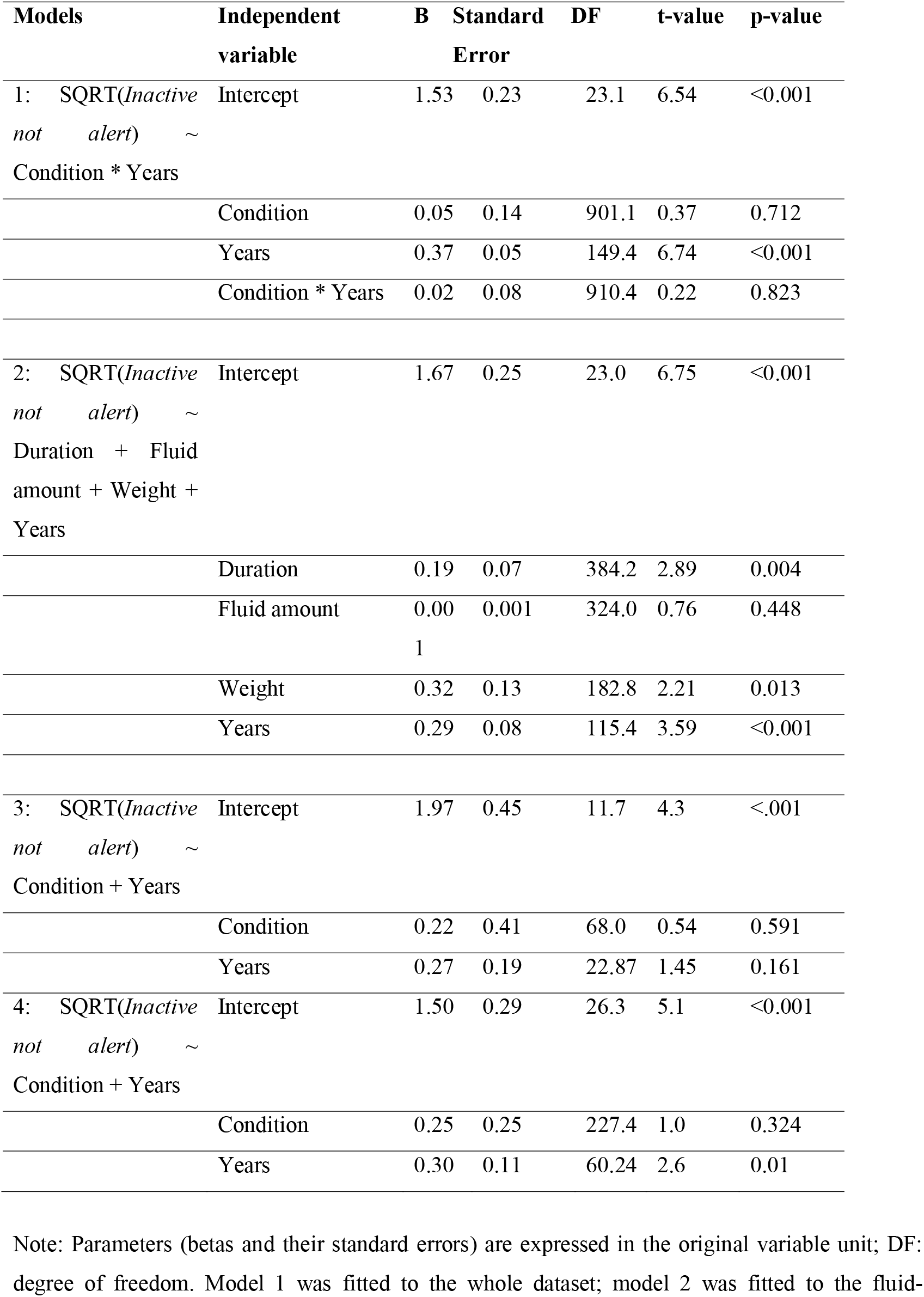
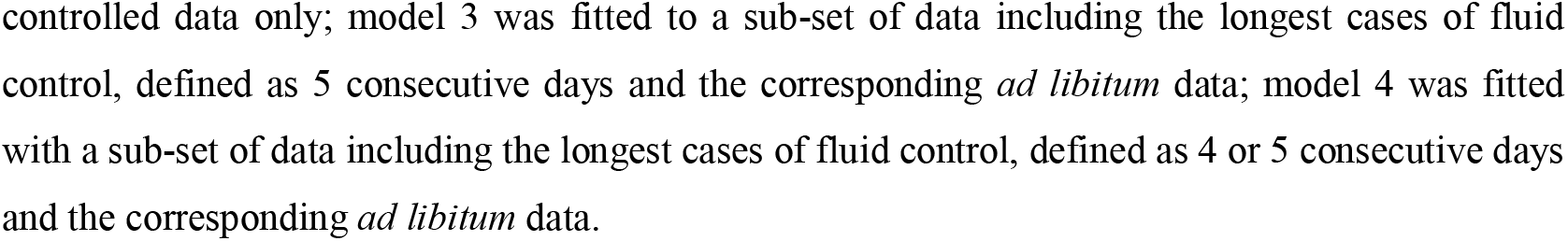
Statistical results for *Inactive not alert* behaviour, after square root transformation.

**Figure 5.**
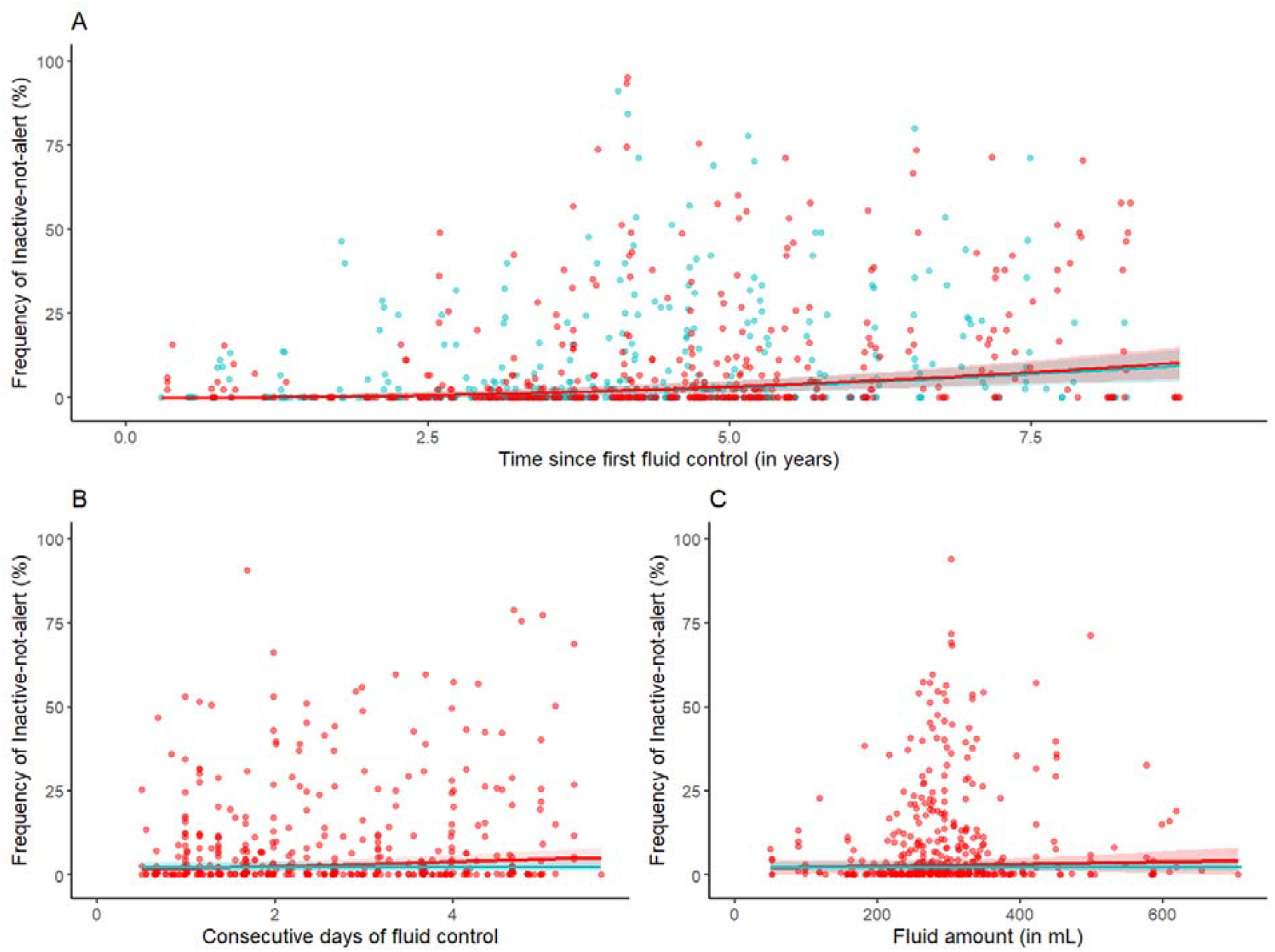
Changes in the frequency of *Inactive not alert* behaviour. (A) Interactive effect of condition (fluid-controlled vs *ad libitum*) with time since first fluid control (in years) (B) Effect of fluid control duration (expressed in number of consecutive days of fluid control). (C) Effect of fluid amount (expressed in ml). Blue: *Ad libitum*; Red: Fluid control. For more details, see legend of Figure 1.

Model 2 revealed a significant effect of duration, with the frequency of *Inactive not alert* increasing with the number of consecutive days of fluid control and no significant effect of fluid amount (Fig. 5B and 5C, Table 4). While the frequency of *Inactive not alert* behaviour increased with duration, the comparison of confidence intervals of the duration effect in model 2 and of the estimated frequency of *Inactive not alert* behaviour when subjects had *ad libitum* access to fluid (i.e. intercept in Model 1) revealed a large overlap, suggesting no significant difference even in the longest instances of fluid control compared to when the animals had *ad libitum* access to fluids (right-hand side of Fig. 5B). This hypothesis was formally tested with model 3, which revealed no significant effect of condition when animals were fluid controlled for 5 consecutive days (Table 4, model 3). Similar results were found if the cases of longest durations were defined as 4 or 5 consecutive days (Table 4, model 4).

## Discussion

This study investigated the effect of fluid control on the psychological well-being of laboratory macaques using a behavioural approach. Results showed no significant acute effect of fluid control (compared to *ad libitum* condition) for any of the behavioural indicators of negative affective state. This absence of significant effect holds not only on average, but also in the most extremes cases of fluid control, namely even after 5 consecutive days of fluid control or when animals received less than 20 ml of fluid per kg. No sensitization (nor habituation) effects were found over years.

Failing to find a significant effect does not equate to proof of no effect. Here we discuss whether the Newcastle fluid control protocol might still induce a negative affective state despite our results; in other words, whether our approach was sensitive enough to detect biologically/psychologically relevant effects.

The choice of a behavioural approach to assess the affective state of the animals was driven by practical constraints. For instance, a cognitive approach such as the judgement bias test, a well-validated alternative approach to assess a wide range of affective states (Bethell, 2015; Lagisz *et al*., 2020; Neville *et al*., 2020), could not be implemented since it is not appropriate for repeated measures with such high frequency (up to 5 days per weeks, during several years). The question to address is therefore whether our behavioural approach might have been insensitive to the negative affective state elicited by fluid control. Our *a priori* hypothesis was that if fluid control had a negative impact on the psychological well-being of macaques, it would induce a high-arousal negative affective state, similar to the effect of thirst in humans (McKinley, 2009). To assess such affective state, we used self-scratching, self-grooming and body shaking. This group of behaviours has been pharmacologically validated in macaques using anxiogenic and anxiolytic drugs (Schino *et al*., 1996), and found to increase when cercopithecines (the primates group to which macaques belong) are exposed to a wide range of stimuli such as exposure to a threatening object (Lacreuse *et al*., 2012) or a human intruder (Raper *et al*., 2013), during agnostic interactions as victim or aggressor (Aureli, Das and Veenema, 1997; Cooper, Aureli and Singh, 2007) and when failing a cognitive task (Leeds and Lukas, 2018), presumably inducing a similarly wide range of high-arousal negative affective states. In addition to these three displacement behaviours, we also included yawning and pacing, despite their lack of validation, on the grounds that they might indicate a distinct negative affective state associated with high or medium arousal. Finally, based on our negative results, we also tested the possibility that contrary to our *a priori* hypothesis, fluid control induced a negative affective state associated with low arousal (rather than high arousal). This was tested by measuring the frequency of *Inactive not alert* behaviour, which has been found to be induced by a wide range of stimuli such as social isolation (Li et al., 2013; Hennessy et al., 2014, 2017), shortening of the photoperiod (Qin *et al*., 2015), and pharmacologically validated using anti-depressant drugs (Perera *et al*., 2011; Qin *et al*., 2015). Considering the wide range of affective states covered by the chosen behavioural indicators, it is therefore unlikely that our lack of significant effects is due to a lack of sensitivity of our approach.

Perhaps the most convincing argument that this interpretation is correct comes from our results of duration and severity. We were able to detect significant changes of displacement behaviour frequency with fluid control severity (defined as fluid amount drunk divided by body weight), and of *Inactive not alert* behaviour frequency with fluid control duration (defined as the number of consecutive days the protocol has been applied at the time of the behavioural measurement). However, it is important to reiterate that the frequency of these behaviours was never significantly higher than during *ad libitum* access to fluid. These results demonstrate that the chosen behaviours were sensitive to variation in affective states elicited by different levels of fluid control severity and duration, but that even in the most extreme cases, the affective state of the animals was not significantly more negative than when animals had *ad libitum* access to fluid.

The final point to consider is whether our lack of significant result is due to a lack of statistical power. Considering that we leveraged an unprecedented amount of data, including 928 observations of 23 individuals (10 females), and applied exclusion criteria designed to increase the sensitivity of the approach (see Material and method section), we think that it is highly unlikely. This interpretation is reinforced by the size of the effects observed in this study. The effect of fluid control on the frequency of displacement behaviours was an increase of 0.5 %. (40.2 vs 40.7 %). For comparison, witnessing an agonistic interaction between conspecifics housed in a different pen, a mild stressor for captive macaques, induces an increase of 24 % in displacement behaviours (Poirier *et al*., 2019) and a low dose of anxiogenic drug provokes an increase of ∼10 % (Schino *et al*., 1996). Considering the intra-individual variation in the frequency of displacement behaviours observed in the present study (full range of frequencies between 0 and 100%, see Figure 1A), an average increase of 0.5% is unlikely to be biologically/psychologically relevant and therefore does not constitute a welfare concern. The day-to-day variation also indicates that other unknown factors are clearly more important in modulating the frequency of displacement behaviours and presumably, the affective states of the animals.

Collectively, these results strongly suggest that the Newcastle fluid control protocol does not compromise the psychological well-being of laboratory macaques. Globally, fluid control protocols vary between research groups (Poirier *et al*., 2021). Our results suggest that some fluid control severity levels and durations might have a negative impact on the psychological well-being of macaques. Neuroscientists using more severe or longer duration of fluid control than that assessed here might want to either refine their protocol or alternatively investigate its psychological impact using the methodology described in this study. It is also important to note that the present dataset prevented us from investigating the effect of fluid control at the individual level, due to a modest sample size for many individuals. As a consequence, we cannot exclude the possibility that the Newcastle protocol had a detrimental impact on the psychological well-being of a minority of individuals. To prevent such risk, we recommend using this protocol in combination with a systematic quantification of displacement and *Inactive not alert* behaviours (behaviours that our study showed to be sensitive to changes in affective states induced by fluid control), in order for researchers to intervene if/when these behaviours become more frequent when an individual is fluid controlled compared to when it is not.

Refining husbandry and experimental procedures is a legal and ethical requirement in the majority of countries using primates in biomedical research (Mitchell *et al*., 2021). To be effective, refinement should be based on scientific evidence. Since our study strongly suggests an absence of adverse effect of the Newcastle fluid control protocol, we argue that when such a protocol is implemented, the welfare of macaques will be better served by focusing resources on refining other procedures.

## Methods

### Subjects and ethical statement

This study re-used data collected for another long-term welfare project on cumulative experience. Twenty-nine laboratory rhesus macaques (*Macaca mulatta*) were initially involved in this project (12 females and 17 males, age range: 3 to 17 years). All individuals expected to be subjected to fluid control during at least two years were included. Animals were bred at UK breeding centres (Medical Research Council’s Centre for Macaques and The Defence Science and Technology Laboratory), where they had been housed with their mother and other adult and juvenile individuals for at least six months (mean weaning age = 1.7 years) and then in groups of juveniles. When adolescents, animals were moved to the Newcastle University primate research facility, which complies with the NC3Rs Guidelines for “Primate Accommodation, Care and Use” (NC3Rs, 2017). In the research facility, animals were socially housed in pairs or trios, in enclosures exceeding the minimal space requirement under the UK legislation (actual size: 2.1 x 3.0 x 2.4 m). Enclosures were enriched with swings, ropes, shelves, novel objects on a biweekly rotation, and had a plain floor covered with wood shavings. Animals were provided with daily foraging opportunities with food spread over the floor, as recommended by LAREF (Reinhardt *et al*., 2007) and the NC3Rs primate welfare guidelines (NC3Rs, 2017). All animals were kept in the same large room, housing over 20 individuals with whom the subjects had visual, olfactory and auditory contact. The facility had natural light, complemented by artificial light based on a 12.5h light/11.5h dark cycle, a humidity rate of 24% and a 20°C temperature.

Animals used in this study were all involved in neuroscience experiments that comprised fluid control and other regulated procedures in accordance with the EU Directive (2010/63/EU), ASPA (1986) and the NIH Guidelines for Care and Use of Animals for Experimental Procedures (National Institutes of Health, 2011), and which were approved by the Home Office for regulated work. The present study, approved by Newcastle University Animal Welfare and Ethical Review Body (project number: ID 928), thus took advantage of fluid control applied for the purpose of the ongoing neuroscience experiments and none of the animals was fluid controlled or kept at Newcastle University for the purpose of the present study.

Each animal was checked daily by technicians, weekly by on-site veterinarians and had an extensive health check every year. Animals’ weights were measured weekly and over the course of the study all animals gained weight in a way appropriate for the species and sex (weight range: 6.2 - 18.3 kg). Animals were considered healthy by the veterinarians and used by neuroscientists as models of healthy human beings.

### Fluid control protocol

The animals were all subjected to the same fluid control protocol. This protocol consisted of a maximum of five consecutive days of fluid control, followed by at least two days of *ad libitum* access to water in the home enclosure. On days of fluid control, experimenters ensured that animals received a minimal amount of fluid. This minimal amount was calculated for each animal, considering their individual water intake when they had *ad libitum* access to water measured during a two-week period. During fluid control days, animals first received fluid as a reward during the experiment. The amount of fluid per experimental trial was controlled by the experimenter but the number of trials was not restricted. In effect, animals could thus get as much fluid as they wanted as long as they worked for it (task difficulty was adjusted to ensure the animals understood the task). At the end of the experimental session, the amount of fluid the animal worked for was measured, and if inferior to the minimal amount the individual was supposed to received, it was supplemented by water. Across the samples, the amount given during fluid control conditions depended on the difficulty of the task and the general motivation of each subject to perform it and varied between 130 and 835 ml/day (see fig 1, panel D for the distribution of the data in ml/kg). On average, animals were fluid controlled for 110 days per year (SD = 69.9; range: 0–290 days). This fluid control protocol is specific to macaques at Newcastle University and may differ significantly from other institutions, where the maximum number of consecutive days of fluid control can be higher, and where there is sometimes no requirement to provide a minimum amount of fluid as long as the animal had the opportunity to earn fluid during the experimental session.

### Indicators of negative affective state

This study re-analysed a pre-existing behavioural database recorded for a different welfare project. This database included the quantification of an exhaustive list of macaque behaviours and a sub-set of behaviours were selected for this study: self-scratching, self-grooming, body shaking, yawning, stereotypic pacing and *Inactive not alert* (Table 5).

**Table 5.** Behavioural ethogram.

| Behavioural item | Description |
| --- | --- |
| Self-grooming | Stroking, picking, or otherwise manipulating own body surface (excluding head post and margins). |
| Self-scratching | Scratching the skin with nails. |
| Body shake | A dog-like shake of the whole body. |
| Yawning | Open the mouth widely, teeth exposed, lips retracted without vocalisation. |
| Pacing | Repetitive (at least twice) walking the same path in the cage with no interruption of more than 2 sec. |
| *Inactive not alert* | Sitting (sometimes with head lower than the shoulders) or lying stationary and alone, while not looking at objects or individuals (eyes may be open or closed), and not doing anything else, for more than 10 seconds. |
| Out-of-sight | The subject is not visible for more than 2 sec. |

### Behavioural recording

Behavioural observations were made using remotely controlled video cameras (Cube HD 1080, Y-cam Solutions Limited, Twickenham, UK, Axis HD Cube and M1065-L, Axis Communications AB, Lund, Sweden) to avoid interference with the animals. The cameras were placed at a distance of 1m -2.5m from the cages, ensuring a broad view of the cage while preventing the cameras from being within reach of the animals. Recording sessions took place in the early morning, starting 15 minutes after the light was switched on in the primate facility (between 6:15 and 7 am), and lasted for 45 minutes. Blue Iris Video Management Software (https://blueirissoftware.com/) was used to automate the recording times. The early morning was chosen as the ideal time for the behavioural data collection as (1) it was the only period during daylight when the focal subjects and their cage-mates were guaranteed to be together in their home cage, as experiments did not start before 8 am; (2) the presence of technicians and researchers was significantly lower than at any other time during daylight hours; and (3) the 15-minute lapse between the light switching on and the start of the data recording ensured that the monkeys were fully awake.

The behavioural data were encoded by 11 human observers blind to the fluid control status of the animals. Observers first underwent extensive video training, followed by a formal assessment where their results were compared to those encoded by an expert observer. For observers who encoded data during more than 6 months, the formal assessment was also repeated several times to avoid any drift in the encoding quality. Inter- and intra-rater reliability was measured using the R.E. coefficient (Maxwell, 1977) and resulted in a mean score of 96.2% (range per session and encoder: 81.9-100%), indicating strong reliability.

### Data pre-processing and statistical analysis

For analysis purposes, the 45 minutes per recording session were divided into 1-minute time bins, in which the presence (1) or absence (0) of a pre-defined list of behaviours was encoded by human observers. Due to the enrichment elements in the cage and the location of the cameras in the facility, the focal subject could not always be seen. When the focal subject was not visible during more than two seconds, the 1-minute time bin was encoded as out-of-sight.

For each recording session, the frequency of each behaviour was calculated as the proportion of 1-minute time bins in which the behaviour was observed, among time bins where the focal individual was not out-of-sight or was seen displaying the behaviour of interest, expressed in percent. The initial dataset comprised data from 1095 sessions from 29 subjects. Further precautions were taken to minimise the probability to get false-negative and false positive results: (1) recording sessions when the focal individual was out-of-sight for 50% or more of the 1-min time bins were discarded (5 sessions, corresponding to 0.5 % of initial dataset); (2) sessions that took place just before or up to 2 days after a general anaesthesia were removed, in case behaviours were influenced by the fasting preceding the procedure or by potential residual lethargy induced by the anaesthesia (20 sessions, 1.8 % of initial dataset); (3) data collected before any fluid control ever happened were discarded (70 sessions, 6.4% of initial dataset) and subjects with less than 1 behavioural session per condition (Fluid controlled and *ad libitum*) at two different time points were excluded (6 subjects, 72 sessions, 6.6 % of initial dataset. After applying these exclusion criteria, the dataset resulted in 928 sessions from 23 subjects (10 females).

Data pre-processing and statistical analyses were performed in R (version 4.4.0, R Core Team, 2023) and RStudio (Posit team, 2025), using the following packages: tidyverse (Wickham *et al*., 2019), lme4 (Bates *et al*., 2015), afex (Singmann *et al*., 2025), ggeffects (Lüdecke, 2018), fitdistrplus (Delignette-Muller and Dutang, 2015), ggpubr (Kassambara, 2023).

This report complies with the ARRIVE guidelines 2.0. Data and R scripts are available <u>here</u>.

## Acknowledgements

This work was funded by a NC3Rs project grant [NC/K000802/1]. J.C.B was supported by a Barbour foundation PhD studentship and A.P. was supported by a UFAW student scholarship. Fluid control data collection was supported by grants to AT [Medical Research Council, UK, G0700976; Wellcome Trust, 093104, and the Biotechnology and Biological Sciences Research Council BB/F021399/1], CIP [Wellcome Trust WT092606AIA, European Research Council Horizon 2020 Consolidator Award, MECHIDENT 724198] and MCC [European Research Council OptoVision 637638, SNSF BSET-0_201532, 310030_197923 and 310030_204544].

## References

Aureli, F., Das, M. and Veenema, H.C. (1997) ‘Differential kinship effect on reconciliation in three species of macaques (Macaca fascicularis, M. fuscata, and M. sylvanus).’, Journal of comparative psychology (Washington, D.C.□: 1983), 111(1), pp. 91–99. Available at: 10.1037/0735-7036.111.1.91.

Bates, D., Mächler, M., Bolker, B. and Walker, S. (2015) ‘Fitting Linear Mixed-Effects Models Using lme4’, Journal of Statistical Software, 67(1), pp. 1–48. Available at: 10.18637/jss.v067.i01.

Bethell, E.J. (2015) ‘A “How-To” Guide for Designing Judgment Bias Studies to Assess Captive Animal Welfare’, Journal of Applied Animal Welfare Science, 18, pp. S18–S42. Available at: 10.1080/10888705.2015.1075833;CSUBTYPE:STRING:SPECIAL;PAGE:STRING:ARTICLE/CHAPTER.

Bleby, J. (1986) ‘Animals (Scientific Procedures) Act 1986.’, The Veterinary record, 119(1), p. 22. Available at: 10.1136/vr.119.1.22-a.

Bonferroni, C. (1936) ‘Teoria statistica delle classi e calcolo delle probabilita’, Pubblicazioni del R Istituto Superiore di Scienze Economiche e Commericiali di Firenze, 8, pp. 3–62.

Cooper, M.A., Aureli, F. and Singh, M. (2007) ‘Sex differences in reconciliation and post-conflict anxiety in bonnet macaques’, Ethology, 113(1), pp. 26–38. Available at: 10.1111/J.1439-0310.2006.01287.X;PAGE:STRING:ARTICLE/CHAPTER.

Delignette-Muller, M.L. and Dutang, C. (2015) ‘fitdistrplus: An R Package for Fitting Distributions’, Journal of Statistical Software, 64(4), pp. 1–34. Available at: 10.18637/jss.v064.i04.

Desimone, R., Olson, C. and Erickson, R. (1992) ‘The Controlled Water Access Paradigm’, ILAR Journal, 34(3), pp. 30–31. Available at: 10.1093/ilar.34.3.30.

European Union (2010) Directive 2010/63/EU of the european parliament and of the council of 22 September 2010 on the protection of animals used for scientific purposes.

Gray, H., Bertrand, H., Mindus, C., Flecknell, P., Rowe, C. and Thiele, A. (2016) ‘Physiological, behavioral, and scientific impact of different fluid control protocols in the rhesus macaque (Macaca mulatta)’, eNeuro, 3(4), pp. 1–15. Available at: 10.1523/ENEURO.0195-16.2016.

Hage, S.R., Ott, T., Eiselt, A.K., Jacob, S.N. and Nieder, A. (2014) ‘Ethograms indicate stable well-being during prolonged training phases in rhesus monkeys used in neurophysiological research’, Laboratory Animals, 48(1), pp. 82–87. Available at: 10.1177/0023677213514043.

Hennessy, M.B., Chun, K. and Capitanio, J.P. (2017) ‘Depressive-like behavior, its sensitization, social buffering, and altered cytokine responses in rhesus macaques moved from outdoor social groups to indoor housing’, Social Neuroscience, 12(1), pp. 65–75. Available at: 10.1080/17470919.2016.1145595.

Hennessy, M.B., McCowan, B., Jiang, J. and Capitanio, J.P. (2014) ‘Depressive-like behavioral response of adult male rhesus monkeys during routine animal husbandry procedure’, Frontiers in Behavioral Neuroscience, 8(September), pp. 1–8. Available at: 10.3389/fnbeh.2014.00309.

Jackowski, A., Perera, T.D., Abdallah, C.G., Garrido, G., Tang, C.Y., Martinez, J., Mathew, S.J., Gorman, J.M., Rosenblum, L.A., Smith, E.L.P., Dwork, A.J., Shungu, D.C., Kaffman, A., Gelernter, J., Coplan, J.D. and Kaufman, J. (2011) ‘Early-life stress, corpus callosum development, hippocampal volumetrics, and anxious behavior in male nonhuman primates’, Psychiatry Research: Neuroimaging, 192(1), pp. 37–44. Available at: 10.1016/j.pscychresns.2010.11.006.

Kassambara, A. (2023) ‘ggpubr: “ggplot2” Based Publication Ready Plots’. Available at: https://CRAN.R-project.org/package=ggpubr.

Lacreuse, A., Gore, H.E., Chang, J. and Kaplan, E.R. (2012) ‘Short-term testosterone manipulations modulate visual recognition memory and some aspects of emotional reactivity in male rhesus monkeys’, Physiology & Behavior, 106(2), pp. 229–237. Available at: 10.1016/J.PHYSBEH.2012.02.008.

Lagarde, D., Laurent, J., Milhaud, C., Andre, E., Aubin, H.J. and Anton, G. (1990) ‘Behavioral effects induced by beta CCE in free or restrained rhesus monkeys (Macaca mulatta)’, Pharmacology Biochemistry and Behavior, 35(3), pp. 713–719. Available at: 10.1016/0091-3057(90)90312-6.

Lagisz, M., Zidar, J., Nakagawa, S., Neville, V., Sorato, E., Paul, E.S., Bateson, M., Mendl, M. and Løvlie, H. (2020) ‘Optimism, pessimism and judgement bias in animals: A systematic review and meta-analysis’, Neuroscience & Biobehavioral Reviews, 118, pp. 3–17. Available at: 10.1016/J.NEUBIOREV.2020.07.012.

Leeds, A. and Lukas, K.E. (2018) ‘Experimentally evaluating the function of self-directed behaviour in two adult mandrills (Mandrillus sphinx)’, Animal Welfare, 27(1), pp. 81–86. Available at: 10.7120/09627286.27.1.081.

Li, X., Xu, F., Xie, L., Ji, Y., Cheng, K., Zhou, Q., Wang, T., Shively, C., Wu, Q., Gong, W., Fang, L., Zhan, Q., Melgiri, N.D. and Xie, P. (2013) ‘Depression-Like Behavioral Phenotypes by Social and Social Plus Visual Isolation in the Adult Female Macaca fascicularis’, PLoS ONE, 8(9). Available at: 10.1371/journal.pone.0073293.

Lüdecke, D. (2018) ‘ggeffects: Tidy Data Frames of Marginal Effects from Regression Models.’, Journal of Open Source Software, 3(26), p. 772. Available at: 10.21105/joss.00772.

Maestripieri, D., Schino, G., Aureli, F. and Troisi, A. (1992) ‘A modest proposal: displacement activities as an indicator of emotions in primates’, Animal Behaviour, 44(5), pp. 967–979. Available at: 10.1016/S0003-3472(05)80592-5.

Major, C.A., Kelly, B.J., Novak, M.A., Davenport, M.D., Stonemetz, K.M. and Meyer, J.S. (2009) ‘The anxiogenic drug FG7142 increases self-injurious behavior in male rhesus monkeys (Macaca mulatta)’, Life Sciences, 85(21–22), pp. 753–758. Available at: 10.1016/j.lfs.2009.10.003.

Maxwell, A.E. (1977) ‘Coefficients of agreement between observers and their interpretation’, British Journal of Psychiatry, 130(1), pp. 79–83. Available at: 10.1192/bjp.130.1.79.

McKinley, M.J. (2009) ‘Thirst’, Handbook of Neuroscience for the Behavioral Sciences [Preprint]. Available at: 10.1002/9780470478509.NEUBB002035.

Mitchell, A.S., Hartig, R., Basso, M.A., Jarrett, W., Kastner, S. and Poirier, C. (2021) ‘International primate neuroscience research regulation, public engagement and transparency opportunities’, NeuroImage, 229, p. 117700. Available at: 10.1016/j.neuroimage.2020.117700.

National Institutes of Health (2011) NIH Guidelines for Care and Use of Animals for Experimental Procedures.

NC3Rs (2017) ‘NC3Rs Guidelines: Non-human primate accommodation, care and use’, National Centre of Replacement, Refinement, & Reduction of Animals in Research, pp. 1–52.

Neville, V., Nakagawa, S., Zidar, J., Paul, E.S., Lagisz, M., Bateson, M., Løvlie, H. and Mendl, M. (2020) ‘Pharmacological manipulations of judgement bias: A systematic review and meta-analysis’, Neuroscience and Biobehavioral Reviews, 108, pp. 269–286. Available at: 10.1016/j.neubiorev.2019.11.008.

Orlans, F.B. (1991) ‘Prolonged Water Deprivation□: A Case Study in Decision Making by an IACUC’, Issues For Institutional Animal Care and Use Committees (lACUCs), 33(3), pp. 48–52. Available at: 10.1093/ilar.33.3.48.

Perera, T.D., Dwork, A.J., Keegan, K.A., Thirumangalakudi, L., Lipira, C.M., Joyce, N., Lange, C., Higley, J.D., Rosoklija, G., Hen, R., Sackeim, H.A. and Coplan, J.D. (2011) ‘Necessity of hippocampal neurogenesis for the therapeutic action of antidepressants in adult Nonhuman primates’, PLoS ONE, 6(4). Available at: 10.1371/journal.pone.0017600.

Pfefferle, D., Plümer, S., Burchardt, L., Treue, S. and Gail, A. (2018) ‘Assessment of stress responses in rhesus macaques (Macaca mulatta) to daily routine procedures in system neuroscience based on salivary cortisol concentrations’, PLoS ONE, 13(1), pp. 1–13. Available at: 10.1371/journal.pone.0190190.

Poirier, C. and Bateson, M. (2017) ‘Pacing stereotypies in laboratory rhesus macaques: Implications for animal welfare and the validity of neuroscientific findings’, Neuroscience and Biobehavioral Reviews, 83, pp. 508–515. Available at: 10.1016/j.neubiorev.2017.09.010.

Poirier, C., Hamed, S. Ben, Garcia-Saldivar, P., Kwok, S.C., Meguerditchian, A., Merchant, H., Rogers, J., Wells, S. and Fox, A.S. (2021) ‘Beyond MRI: on the scientific value of combining non-human primate neuroimaging with metadata’, NeuroImage, 228. Available at: 10.1016/j.neuroimage.2020.117679.

Poirier, C., Oliver, C.J., Castellano Bueno, J., Flecknell, P. and Bateson, M. (2019) ‘Pacing behaviour in laboratory macaques is an unreliable indicator of acute stress’, Scientific Reports, 9(1). Available at: 10.1038/s41598-019-43695-5.

van de Pol, M. and Wright, J. (2009) ‘A simple method for distinguishing within-versus between-subject effects using mixed models’, Animal Behaviour, 77(3), pp. 753–758. Available at: 10.1016/J.ANBEHAV.2008.11.006.

Posit team (2025) ‘RStudio: Integrated Development Environment for R’. Boston, MA. Available at: https://www.posit.co.

Prescott, M.J., Brown, V.J., Flecknell, P.A., Gaffan, D., Garrod, K., Lemon, R.N., Parker, A.J., Ryder, K., Schultz, W., Scott, L., Watson, J. and Whitfield, L. (2010) ‘Refinement of the use of food and fluid control as motivational tools for macaques used in behavioural neuroscience research: Report of a Working Group of the NC3Rs’, Journal of Neuroscience Methods, 193(2), pp. 167–188. Available at: 10.1016/j.jneumeth.2010.09.003.

Qin, D., Chu, X., Feng, X., Li, Z., Yang, S., Lü, L., Yang, Q., Pan, L., Yin, Y., Li, J., Xu, L., Chen, L. and Hu, X. (2015) ‘The first observation of seasonal affective disorder symptoms in Rhesus macaque’, Behavioural Brain Research, 292(32), pp. 463–469. Available at: 10.1016/j.bbr.2015.07.005.

R Core Team (2023) ‘R: A Language and Environment for Statistical Computing’. Vienna, Austria. Available at: https://www.R-project.org/.

Ralph, C.R. and Tilbrook, A.J. (2016) ‘Invited Review: The usefulness of measuring glucocorticoids for assessing animal welfare’, Journal of Animal Science, 94(2), pp. 457–470. Available at: 10.2527/jas.2015-9645.

Raper, J., Wilson, M., Sanchez, M., Machado, C.J. and Bachevalier, J. (2013) ‘Pervasive alterations of emotional and neuroendocrine responses to an acute stressor after neonatal amygdala lesions in rhesus monkeys’, Psychoneuroendocrinology, 38(7), pp. 1021–1035. Available at: 10.1016/J.PSYNEUEN.2012.10.008,.

Reinhardt, V., Rodgers, J.C., Shayne, K., Schwartz, J., Bell, L., Vila, P.M., Kerwin, A., Rodgers, J.C., Schultz, P., (Genny), G.A.-K., Donnelly, M., Augusto, V., Moore, A. and Cawston, D. (2007) A Discussion Relating to Animals in the Research La.

Schino, G., Perretta, G., Taglioni, A.M., Monaco, V. and Troisi, A. (1996) ‘Primate displacement activities as an ethopharmacological model of anxiety’, Anxiety, 2(4), pp. 186–191. Available at: 10.1002/(SICI)1522-7154(1996)2:4<186::AID-ANXI5>3.0.CO;2-M.

Singmann, H., Bolker, B., Westfall, J., Aust, F. and Ben-Shachar, M.S. (2025) ‘afex: Analysis of Factorial Experiments’. Available at: https://CRAN.R-project.org/package=afex.

Wegener, D., Oh, D.Q.P., Lukaß, H., Böer, M. and Kreiter, A.K. (2021) ‘Blood analysis of laboratory macaca mulatta used for neuroscience research: Investigation of long-term and cumulative effects of implants, fluid control, and laboratory procedures’, eNeuro, 8(5). Available at: 10.1523/ENEURO.0284-21.2021.

Westlund, K. (2012) ‘Can conditioned reinforcers and Variable-Ratio Schedules make food- and fluid control redundant? A comment on the NC3Rs Working Group’s report’, Journal of Neuroscience Methods, 204(1), pp. 202–205. Available at: 10.1016/j.jneumeth.2011.03.021.

Wickham, H., Averick, M., Bryan, J., Chang, W., McGowan, L.D., François, R., Grolemund, G., Hayes, A., Henry, L., Hester, J., Kuhn, M., Pedersen, T.L., Miller, E., Bache, S.M., Müller, K., Ooms, J., Robinson, D., Seidel, D.P., Spinu, V., Takahashi, K., Vaughan, D., Wilke, C., Woo, K. and Yutani, H. (2019) ‘Welcome to the tidyverse’, Journal of Open Source Software, 4(43), p. 1686. Available at: 10.21105/joss.01686.

Yamada, H., Louie, K. and Glimcher, P.W. (2010) ‘Controlled Water Intake: A Method for Objectively Evaluating Thirst and Hydration State in Monkeys by the Measurement of Blood Osmolality’, J Neurosci Methods, 191(1), pp. 83–89. Available at: 10.1016/j.jneumeth.2010.06.011.Controlled.

